# Sleep after Motor Sequence Learning Enhances Post-Movement Parietal Beta Synchronization

**DOI:** 10.1101/2025.10.16.682965

**Authors:** Marie-Frédérique Bernier, Roxane S. Hoyer, Françoise Lecaignard, Alain Nicolas, Olivier Bertrand, Philippe Albouy, Geneviève Albouy

**Affiliations:** CERVO Brain Research Center, Québec, Canada; School of Psychology, Faculty of social sciences, Laval University, Québec, Canada; Lyon Neuroscience Research Center (CRNL) - Lyon 1 University, Inserm U1028, CNRS UMR5292, Lyon, France; International Laboratory for Brain, Music and Sound Research (BRAMS), CRBLM, Montreal, Canada; CHU de Québec research center, Université Laval, Québec, QC, Canada; Department of Health and Kinesiology, University of Utah, Salt Lake City, USA

## Abstract

The neural substrates supporting the beneficial effect of sleep on motor memory consolidation are well described. However, less is known about the brain oscillatory dynamics underlying these processes. We characterized the oscillatory dynamics associated with motor sequence learning and their modulation by post-learning sleep using magnetoencephalography (MEG) in young healthy adults. After learning a motor sequence task while their brain activity was recorded with MEG, participants were distributed in two groups according to whether they slept or were totally sleep deprived during the first post-training night. Consolidation was assessed with a retest in the MEG three days after training. Behaviorally, performance improved over the consolidation interval irrespective of whether sleep was afforded during the first night. MEG results showed that initial motor sequence learning was characterized by a progressive decrease in beta Event Related Desynchronization (ERD, 18-25Hz) over bilateral motor areas. Interestingly, while these practice-related modulations of beta ERD were not influenced by the sleep status, post-learned-movement beta Event Related Synchronization (ERS) over bilateral parietal areas increased over the consolidation interval in the sleep, compared to the sleep deprived, group. These results extend current models of motor memory consolidation by identifying ERS as an oscillatory marker of sleep-dependent consolidation.

## Introduction

Acquiring new motor skills is inherently part of our everyday life and allows us to learn to execute complex movements like hitting a tennis ball, quickly typing on a keyboard, or taking a sip of coffee. A large majority of these motor skills consists of sequences of movements that are learned through repeated practice (Doyon et al. 2009). During the acquisition of a new skill in procedural memory, performance rapidly improves online during task practice, but slower performance improvements are also observed after offline periods during which the task is not practiced (Karni et al. 1995; Korman et al. 2003; Korman et al. 2007). These offline periods between practice sessions are thought to be critical windows during which memory consolidation, the process by which new memories are reorganized into stable ones, occurs (Robertson et al. 2004). And there is consistent evidence in the literature that sleep plays a critical role in this offline motor memory consolidation process (see King et al. 2017 for review).

The cerebral processes supporting online motor sequence learning and sleep-related offline motor memory consolidation have been extensively characterized using functional magnetic resonance imaging (fMRI). Earlier research indicates that cerebello-striato-cortical and hippocampo-cortical networks are critically involved in motor learning and memory consolidation (see Albouy et al. 2013a; Doyon et al. 2009; King et al. 2017; Penhune and Steele 2012 for reviews). However, less is known about the brain *oscillatory dynamics* related to these processes. Earlier electrophysiological research has shown that preparation, execution, and imagination of a movement produce an event-related desynchronization (ERD), i.e. a decrease in the magnitude of brain oscillations as compared to baseline, over sensorimotor areas in the beta (14-25 Hz) band (Pfurtscheller and Lopes da Silva 1999; Neuper et al. 2006, Manganotti et al. 1998). Interestingly, this electrophysiological marker is subject to plasticity processes as later stages of learning are accompanied by decreased movement-related ERD in the beta band on motor regions contralateral to the movement performed (Formaggio et al. 2008; Pineda 2005; Dyck and Klaes 2024). In addition to ERD, post-movement (rest) periods are characterized by event-related synchronization (ERS) processes, i.e. an increase in the magnitude of beta oscillatory activity as compared to baseline over the brain regions involved during task practice (Jurkiewicz et al. 2006; Pfurtscheller and Lopes da Silva 1999; Neuper et al. 2006). This phenomenon has been associated with active motor inhibition during rest periods following movements (Heinrichs-Graham et al. 2017), and more recently, with rapid motor memory consolidation (Bönstrup et al. 2019, 2020; Buch et al. 2021). Specifically, post-movement brain activity observed over frontoparietal network during rest blocks interspersed with task practice during initial motor sequence learning is thought to reflect the reactivation of task-related brain patterns that support a rapid form of motor memory consolidation (Bönstrup et al. 2019; Buch et al. 2021). Importantly, it is currently unknown how post-learning sleep influences these oscillatory markers of initial motor sequence learning.

The goal of this study was therefore to characterize the oscillatory correlates of motor sequence learning and examine the effect of sleep on these learning-related markers. To do so, we used MEG to record brain activity of young healthy participants while they learned a 5-element sequential finger-tapping task with their left non-dominant hand. After learning, participants were divided in two groups according to whether they slept normally (SG) or were totally sleep-deprived (SDG) during the first post-training night. Brain activity was then recorded with MEG during a retest session 72h after initial learning to assess the effect of sleep on the electrophysiological correlates of motor memory consolidation. Source-localized beta-band oscillatory activity (18-25 Hz) was extracted from the whole brain for both practice and inter-practice rest periods during the training and retest sessions. According to studies showing that sleep favors motor memory consolidation (see King et al. 2017 for a review), we hypothesized that changes in motor performance between the training and the retest sessions would be greater in the SG as compared to the SDG. Based on evidence that later stages of learning are characterized by decreased movement-related ERD in the beta band on the motor-cortical electrodes (Formaggio et al. 2008; Pineda 2005; Dyck and Klaes 2024), we hypothesized that desynchronization would decrease as a function of learning in the beta band over motor and premotor regions. In the same vein, we expected to observe decreased desynchronization after a consolidation period and hypothesized that sleep – as compared to sleep deprivation – would further potentiate this effect. Last, based on studies showing that the rapid form of consolidation occurring during post-movement rest blocks is particularly pronounced early during initial learning and supported by modulations of beta power in a frontoparietal and sensorimotor-hippocampal networks (Bönstrup et al. 2019; Buch et al. 2021), we expected to observe a decrease in the post-movement beta ERS magnitude throughout practice in sensorimotor and parietal regions. As this form of consolidation occurs early during learning, we did not expect sleep - as compared to sleep deprivation - to modulate these effects.

## Material and Methods

### Ethics

All procedure were approved by the local Ethics Committee on Human Research, the Ethics Committee of Protection of Persons *Sud-Est II* in Lyon (ethics committee approval number DGS2006-0101). Note that data analysis at Laval University has been approved by the CERUL of Université Laval (2022-431/14- 11-2022).

### Participants

MEG recordings were obtained from 35 healthy participants at the MEG department of the CERMEP-Imagerie du Vivant (Lyon, France). Participants were randomly distributed in two groups that included 19 and 16 participants in the sleep group (SG) and sleep-deprived group (SDG), respectively. Note that 2 participants were discarded from the analyses. One participant (SG) was excluded because he was outlier in electrophysiological responses and performance (with the mean amplitude of cerebral responses during training above the mean plus 2 standard deviations computed on the population, and another participant (SG) was excluded from the analyses because they did not respect the constant sleep schedule between the training and retest sessions. Eventually, 33 participants (mean age: 22.7±3.6 years) were included for the analyses, 17 in the SG and 16 participants in the SDG. All participants were right-handed (Edinburgh Handedness Questionnaire; Oldfield 1971), self-reported no current or previous history of neurological, psychological, or psychiatric conditions, and were free of any psychotropic or sleep impacting medications. None of the participants worked night shifts or took trans-meridian trips in the two months preceding the experiment. None of the participants received previous extensive training as a musician or as a professional typist. Further screening using standardized questionnaires also ensured that participants did not exhibit any indication of anxiety (Beck Anxiety Inventory (BAI); Beck et al. 1988) or depression (Beck Depression Inventory (BDI); Beck et al. 1961). Participants reported normal sleep quality and quantity during the month prior to the study (Pittsburgh Sleep Quality Index (PSQI); Buysse et al. 1989) as well as normal levels of daytime sleepiness (Epworth Sleepiness Scale; Johns 1991). None of the participants were extreme evening or morning type individuals (Morningness-Eveningness Questionnaire (MEQ); Horne & Östberg 1976). Participants received a monetary compensation and gave their written informed consent to participate in the study.

### Experimental procedure

Three days before the first experimental session, participants were instructed to follow a constant sleep schedule (according to their own rhythm *±* 1 hour) and to keep the same schedule for three more days until their second visit for the retest session. Compliance with the schedule was assessed using sleep diaries and wrist actigraphy (Cambridge Neuroscience, Cambridge, UK). On the first experimental day, participants were trained on an explicitly known 5-element finger sequence (Figure 1A) in the MEG. After the training session, the participants were randomly assigned to one of two groups according to whether they would be afforded regular sleep during the three post-training nights (Sleep Group: SG) or be totally sleep-deprived for the first night following initial training (Sleep Deprived Group: SDG). They were informed of their group assignment after the first training session (Figure 1B). SG participants went back home after the first training session and were instructed to continue respecting the regular sleep schedule monitored with sleep diary and wrist actigraphy. Participants in the SDG stayed awake during the entire post-training night at the Hypnological Exploration Unit – University Hospital Service of Psychiatry in the Hospital Centre Le Vinatier (Lyon, France) and were constantly monitored by an experimenter. During the sleep deprivation night, the lights were dimmed, and physical activity was maintained as minimal as possible. Participants were able to get up each hour and eat a snack. After the night of sleep deprivation, participants were instructed to not sleep the next day and perform their habitual activities. The retest session took place in the MEG 72 hours after the training session for both groups, allowing two nights of recovery sleep for the SDG. On experimental day 4, the retest session was administered at the same time of the day as the training session (ranging between 8 am to 7 pm across participants), to account for possible fluctuations in performance due to the circadian rhythm.

**Figure 1.**
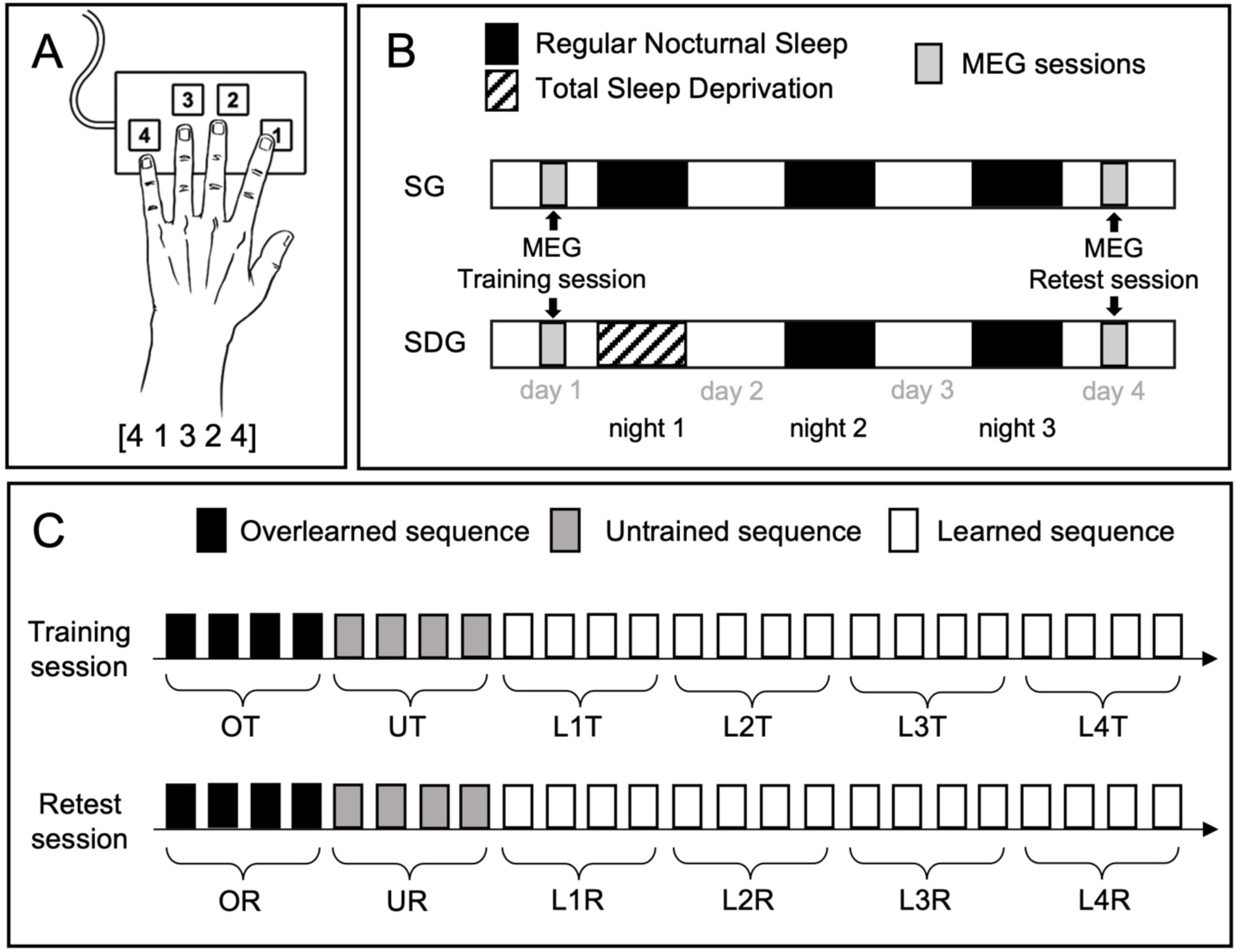
Task and experimental design. **A:** Finger Tapping Task, FTT. Participants performed an explicitly known 5-element finger sequence as rapidly and accurately as possible with their left non-dominant hand on a keyboard within the MEG. **B:** Experimental design. Participants’ brain activity was recorded with MEG during the training session, and they were then divided into 2 groups according to their sleep condition on the first post-training night (SG: Sleep Group, SDG: Sleep Deprived Group). All participants were retested in the MEG three days later. **C:** Task design. Training (T) and retest (R) sessions consisted of 24 30s-practice blocks separated by 15s-rest blocks on different conditions (overlearned (O), untrained (U) and learned (L) sequences). Curly brackets represent the way blocks were collapsed for the MEG analyses (see methods).

### Task

#### Behavioral data acquisition

Participants performed a motor sequence learning task on a specialized MEG-compatible keyboard in the MEG on two different occasions referred to as the Training and the Retest sessions. The finger tapping task (FTT, Figure 1A) was coded in Presentation Version 11.0 (http://www.neurobs.com) and required participants to tap on a keyboard, with their left non-dominant hand, a 5-element finger sequence as rapidly and accurately as possible (Figure 1A). The sequence was explicitly known by the participants and was continuously presented on the screen during task practice (but not during rest blocks). During rest periods, a red cross was presented on the screen, and the participants had the instructions to fixate the red crossed and keep their fingers still on the keyboard. Three types of sequences were practiced: Overlearned O (4 3 2 1, where 1 corresponds to the left index finger and 4 to the left pinky finger), Untrained U (U1: 4 2 3 1 4 or U2: 1 4 2 3 1 or U3: 3 1 4 2 3 or U4: 2 3 1 4 2, order counterbalanced across sessions) and Learned L (4 1 3 2 4) (Figure 1C). The O and U conditions were designed to control for speed/complexity and learning respectively, whereas the L condition was the sequence learned through repetitive practice. The Training and Retest sessions were identical and included 24 practice blocks of 30 seconds each separated by rest periods of 15 seconds. The two sessions consisted of 4 blocks of the O sequence, 4 blocks of the U sequence and 16 blocks of the L sequence.

#### Behavioral data analyses

Similar as in our previous research (Albouy et al. 2013b, Albouy et al. 2015), the primary outcome variable consisted in the speed measure, corresponding to the time (ms) needed to perform a chunk of 3 correct button-presses (chunk of 3 correct responses were used in the behavioral analysis to match the events used in the MEG analyses, see below). As accuracy remained high and did not change as a function of learning, corresponding resulted are reported in the supplements. Given that the speed data did not follow a normal distribution, a log transformation was applied on the speed data. Behavioral data were analyzed with repeated-measure analyses of variance (ANOVA). Specifically, performance on the different task conditions (O, U and L) and the different sessions (training and retest) were analyzed with separate ANOVAS using block as within-participant factor (4 or 16 for O/U and L conditions, respectively) and group (SG and SDG) as between-participant factor. Note that one participant (SG) was excluded from the analyses of the U condition as this participant practiced a wrong sequence during the second block of UR. A separate ANOVA using the four last blocks of the L sequence as a within-participants factor and group (SG vs. SDG) as a between-participants factor was performed to test whether performance reached plateau levels before the consolidation interval in a similar way between the 2 groups. To test whether changes in performance between training and retest differ between groups, performance was first averaged over the last four blocks of training and over the four first blocks of retest separately for each condition and compared (i.e., OT vs. OR for the O condition, UT vs. UR for the U condition and L4T vs. L1R for the L condition) between groups using an ANOVA with session (Training and Retest) as within-participant factor and group (SG and SDG) as between-participant factor. Analyses were performed in R using the emmeans package (emmeans version 1.6.3). The results of the main analyses were considered significant at *p* < .05. For significant main effects or interactions, post hoc tests were systematically performed using the emmeans package (emmeans version 1.6.3). *P-*values were considered significant at *p* < .05 and were adjusted for the number of performed comparisons (Tukey).

### MEG

#### MEG data acquisition

Recordings were carried out using a whole-head, 275-channel MEG system (CTF-275 by VSM MedTech Inc., Vancouver, Canada) with continuous sampling at a rate of 600 Hz, a 0–150 Hz filter bandwidth, and first-order spatial gradient noise cancellation. Electromyography (EMG) was recorded from a bipolar montage of two surfaces electrodes overlying the abductor muscles of the fingers on the left forearm to allow the pressure of the fingers on the buttons to be recorded continuously for each finger. Electrocardiography (ECG) was recorded from a bipolar montage of two electrodes placed on the chest. Head position was determined with coils fixated at the nasion and the preauricular points (fiducial points). Head position was acquired continuously (continuous sampling at a rate of 150 Hz) and checked at the beginning of each block to ensure that head movements did not exceed 0.5 cm (this was confirmed by additional offline checking before the data analyses). The positions of the fiducial coils were measured relative to the participant’s head using a 3-D digitizer system (Polhemus Isotrack). To facilitate anatomical registration with MRI, about 150 additional scalp points were also digitized. The participants were seated upright in the magnetically shielded recording room and practiced the sequence on a MEG-compatible, optic-fiber, 4-digit keyboard.

#### MEG preprocessing

MEG data were preprocessed using Brainstorm (3.4.0.0 version), a software package for analyzing electrophysiological data (Tadel et al. 2011). The contamination artefacts that interfere with the data analysis, such as blinking and eye movements (saccades), cardiac activity, and environmental noises, were reduced with signal space projection (SSP). Heartbeat and eye blink events were automatically detected on the ECG, and the frontal sensor MLF14 (as no EOG was recorded), respectively. Projectors were defined using principal component analysis (PCA) of these data segments filtered between 10 and 40 Hz (for heartbeats) or 1.5 and 15 Hz (for eye blinks) in a 160 ms time window centred around the heartbeat event, or 400 ms around the eye blink event (Brainstorm default parameters settings). The principal components that best captured the artifact’s sensor topography were manually selected as the dimension against which the data was orthogonally projected away.

The first principal component was sufficient for most participants to remove artifact contamination. The projectors obtained for each participant were propagated to the corresponding MEG source imaging operator. A notch filter was applied to remove the main frequency and the harmonics of the electrical signal from the wall outlets (50, 100 and 150 Hz), and a band-pass filter between 0.3 and 80 Hz was applied. The events were then imported in chunks of 3 correct button presses (i.e., each button press followed by two successive correct button presses), and data were epoched between -1000 ms to 2000 ms relative to the onset of the first button press of the chunk. Data of the inter-practice rest periods were epoched from -5000 to 20000 ms relative to the beginning of the rest period. The trials exceeding ± 3000fT at any sensor during an event time window were rejected. A noise covariance matrix was computed using a 10-minutes empty room recording for each session and head modelling were performed using overlapping spheres and MNI templates.

#### MEG data analyses

Brain oscillatory activity was localized with a source reconstruction of the raw data on the cortical surface using standard brain MRIs. Source reconstruction was performed using tools available in Brainstorm, all with default parameter settings (Tadel et al. 2011). Forward modeling of neural magnetic fields was performed using the overlapping-sphere technique (Huang et al. 1999). The lead fields were computed from elementary current dipoles distributed perpendicularly to the cortical surface of each individual (Baillet et al. 2001). MEG source imaging was performed by linearly applying Brainstorm’s weighted-minimum norm operator onto the preprocessed data. The data were previously projected away from the spatial components of artifact contaminants. For consistency between the projected data and the model of their generation by cortical sources, the forward operator was projected away from the same contaminants using the same projector as the MEG data.

Hilbert Transforms were computed on the source reconstructed data, for each correct chunk during practice and for each 15-second rest periods, of the training and retest sessions. We focused on Event Related Desynchronization (ERD) during task practice and Event Related Synchronization (ERS) during inter-practice periods in the beta range (18-25 Hz) as markers of motor learning (Pfurtscheller 2001). For task periods, the magnitude of beta activity between 18 and 25 Hz was extracted for each correct chunk of the conditions (O, U and L) and all vertices, and was z-scored using the average activity of the chunk time window (-400 to 1400 ms, a time period that does not include edge effects associated with the Hilbert transform). This entire window was used as the baseline for z-scoring because button presses were too close during task performance to allow for a pre-button press baseline (which would contain activity associated with the preceding button press). This analysis was performed for all conditions, and Hilbert data were then averaged across 4 blocks of practice for each condition and for each participant (OT, UT, L1T, L2T, L3T, L4T, OR, UR, L1R, L2R, L3R and L4R, see Figure 1C). The mean beta magnitude for these blocks were extracted for each participant to contrast conditions, sessions and groups. For the rest periods, the magnitude of beta activity between 18 and 25 Hz was extracted for each 15-seconds inter-tapping rest periods for all conditions (OTrest, UTrest, L1Trest, L2Trest, L3Trest, L4Trest, ORrest, URrest, L1Rrest, L2Rrest, L3Rrest and L4Rrest) and all vertices, Hilbert maps were z-scored by the practice activity between -4000 to 0 ms relative to the beginning of the rest period. The mean beta magnitude for these rest periods were extracted for each participant to contrast conditions, sessions and groups.

#### MEG statistical analyses

MEG contrasts were performed at the whole brain level with non-parametric paired tests using cluster-level statistics (as implemented in Fieldtrip, Oostenveld et al. 2011) on the source maps. The tests were performed on data averaged over time and clusters were based on spatial adjacencies between vertices at the source level (see below for a description of the data-driven approach used to determine time-window over which effects were averaged). The analyses of MEG data were performed using Brainstorm (3.4.0.0 version, Tadel et al. 2011) and MATLAB (9.9 version).

#### Definition of time periods of interest

To define time windows in which ERD and ERS fluctuations were maximal at the whole brain level for both practice and inter-practice rest periods, we performed clustered based permutation tests corrected in time and space across all conditions on the training data. Specifically, for ERD, we used the training data collapsed across all conditions to compare oscillatory activity between a 0 to 500ms post-button-press window and 500 ms baseline window extracted from an independent resting state recording acquired before the first training session. For ERS, we contrasted the oscillatory activity between 2000 to 14000 ms and - 4000 to 0 ms baseline window relative to the beginning of the rest period for inter-practice rest periods. All statistical analyses presented below were then performed at a whole-brain level using the time average of the significant time windows identified with this analysis (see results section and Figure 3A). Even though all the analyses contrasting the different conditions and sessions were performed at the whole brain level, beta values were extracted – for illustration purposes – from spatial clusters identified in this first analysis showing significant ERD and ERS modulation across all conditions of the training session. Note that the whole brain statistical maps are also shown for completeness. The practice and rest spatial clusters used for beta extraction were defined using a 15% threshold around the highest peak of activity in the most significant spatial cluster (see Figure 3A).

#### Within session (training, retest) contrasts

For the training and retest sessions, separately, we first investigated whether practice-related ERDs were modulated by task condition and learning. Using cluster-based permutation tests, we first contrasted data (averaged over time, see above) from the U and O conditions within each session separately (i.e., UT vs. OT and UR vs. OR) to assess the effect of task difficulty and performance speed. We then investigated the main effect of learning within each session by comparing the electrophysiological responses of the L and U conditions (i.e., UT vs. L1T, L2T, L3T, L4T and UR vs. L1R, L2R, L3R, L4R). We also compared the different learning stages (associated with different speed performance) with the overlearned condition (i.e., OT vs. L1T, L2T, L3T, L4T and OR vs. L1R, L2R, L3R, L4R). Finally, practice-related changes in brain activity were evaluated by comparing the different L blocks (L1, L2, L3, L4) with one another within each session. Group comparisons (SG vs. SDG) were performed on the retest data to evaluate the effect of sleep on the contrasts described above.

#### Between sessions contrasts

Changes in oscillatory activity between training and retest were evaluated within each group and between groups by comparing whole brain beta magnitude of the first four blocks of the L condition during retest (L1R) with the four last blocks of the L condition during training (L4T) for practice and rest (L1R vs. L4T). Those blocks were used to match the behavioral analyses examining offline changes in performance over the consolidation interval. Between-session changes in oscillatory activity were also examined for the O and U conditions.

#### Correlations

Correlation analyses were performed to examine the relationship between the learning- and sleep-related modulation in beta power and the behavioral marker of consolidation (i.e., offline changes in performance speed and accuracy between training and retest). The brain imaging contrast used for these correlations analyses are the following: L1T vs. L4T practice, L1R vs. L4R rest and L4T vs. L1R rest. These analyses were performed with whole brain Pearson’s correlation, FDR corrected p< .05.

## Results

### Behavior

We first aimed to identify practice-related behavioral changes during the training session. The repeated-measures ANOVAs performed on speed during training (using block as a within-subject factor and group as between participant factor) showed a main effect of block for the overlearned (O) and learned (L) sequences (O, F(3,124)=4.465, *p* < .01, η^2^=.09 and L, F(15,496)=6.877, *p* < .001, η^2^=.17) whereby time to perform a correct chunk of 3 elements decreased as a function of practice. No such effect was observed for the untrained (U) sequence condition (U, F(3,120)=1.424, *p* = .24, η^2^=.03) as a new sequence was introduced in each block. As expected, there was no significant effect of group for the U and L conditions (U, F(1,120)=0.958, η^2^=.008; L, F(1,496)=0.319, η^2^=.0005; all *ps* > .3), but a group effect was observed for the O condition (O, F(1,124)=10.287, p < .01, η^2^=.07), where participants in the SDG were initially faster than those in the SG on this particular condition. The block by group interaction was not significant for any of the conditions (O, F(3,124)=0.167, η^2^=.003; U, F(3,120)=0.103, η^2^=.002; L, F(15,496)=0.092, η^2^=.002; all *ps* >.9) indicating that performance similarly improved in both groups on the different conditions during training (Figure 2A). Next, we examined whether performance on the learned sequence reached a plateau at the end of training. This effect was tested with an ANOVA using the four last blocks of the L sequence as a within-participants factor and group (SG vs. SDG) as a between-participants factor. The analysis revealed that the block effect was non-significant (F(3,124)=0.016, *p* > .9, η^2^=.0004), and neither a group effect (F(1,124)=0.044, *p* > .8, η^2^=.0004) nor a group by block interaction (F(3,124)=0.114, *p* > .9, η^2^=.003) were observed. These results suggest that performance did plateau at the end of training in a similar way between groups (Figure 2A).

**Figure 2.**
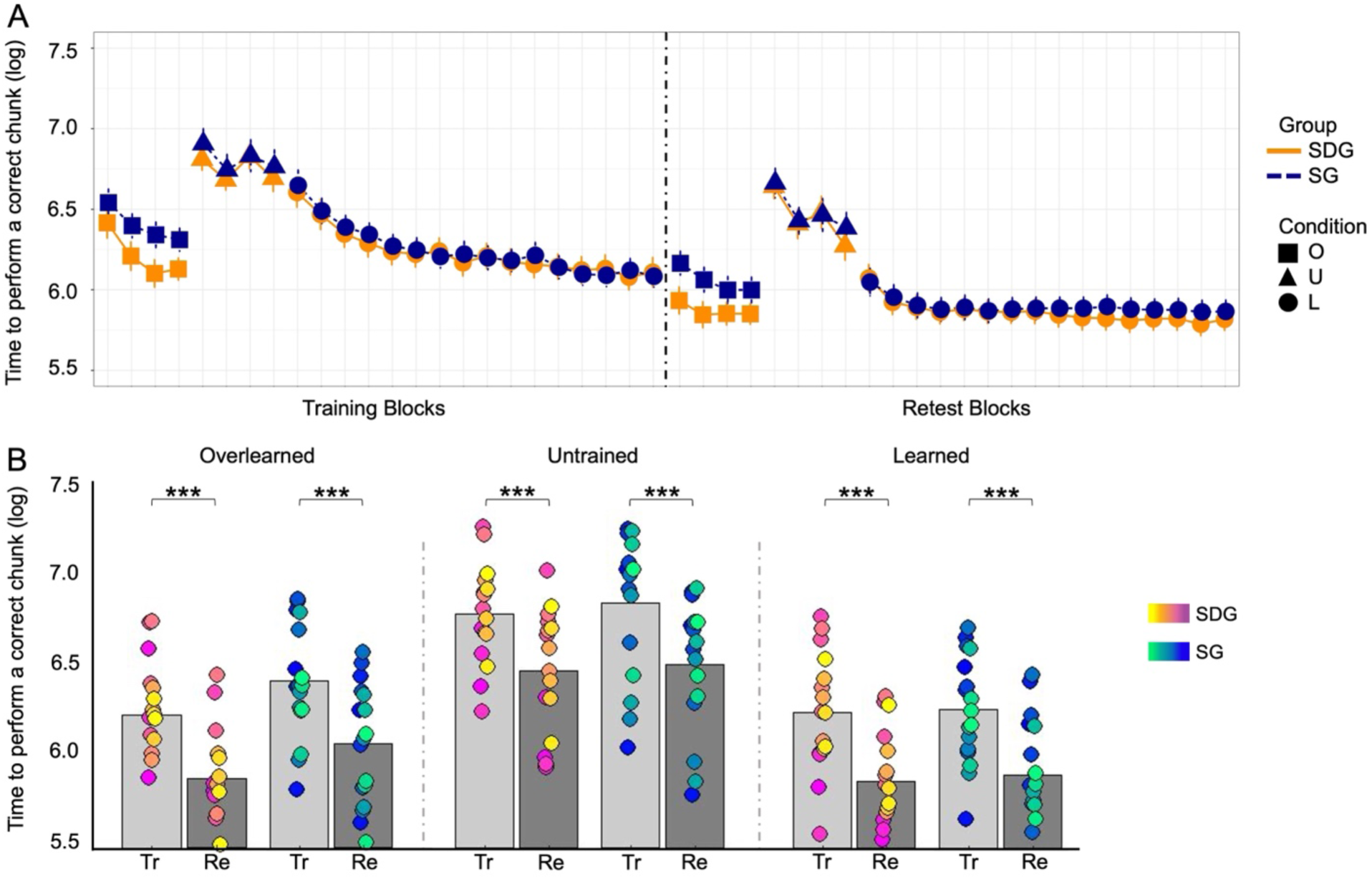
Behavioral results (speed). **A:** Performance speed (time to perform a chunk of 3 correct button- press in ms) for each block of each condition during training and retest sessions for participants in the sleep group (SG, blue) and the sleep-deprived group (SDG, orange). Error bars represent SEM. **B:** Average performance across the last 4 blocks of training (Tr) and the first 4 blocks of retest (Re) for each condition in both groups (SDG, spring color bar; SG, winter color bar). Circles indicate individual participants. Significant (*) offline gains in performance were observed in all conditions and in both groups.

We then aimed to identify whether performance during retest was influenced by the sleep status of the first post-training night. The repeated-measures ANOVAs performed on performance speed during retest (using block as a within-subject factor and group as between participant factor) showed a main effect of block for the U sequence (U, F(3,120)=4.468, *p* < .01, η^2^=.1), but not for the O and L sequences (O, F(3,124)=1.187, η^2^=.03; L, F(15,496)=1.120, η^2^=.03, all *ps* > .3). In contrast to our expectations, we did not observe any significant effect of group for the U and L conditions (U, F(1,120)=0.281, η^2^=.0.002; L, F(1,496)=1.912, η^2^=.004 all *ps* > .1). However, similar as during the training session, we observed a significant effect of group for the O sequence (O, F(1,124)=11.678, *p* < .001, η^2^=.08), whereby performance was faster in the SDG as compared to the SG. The block by group interaction was not significant for any of the conditions (O, F(3,124)=0.169, η^2^=.004; U, F(3,120)=0.175, η^2^=.004; L, F(15,496)=0.077, η^2^=.002, all *ps* > .9) indicating that block-to-block changes in performance during retest were similar in both groups on the different conditions (Figure 2A). Altogether, these results suggest that sleep during the first post-training night did not influence overall performance at the 72h retest.

We then tested the effect of sleep (deprivation) on between-session changes in performance, reflecting offline consolidation processes, for each condition using an ANOVA with session (i.e., the average of the last four blocks of the training session against the first four blocks of the retest session) as a within-participants factor and group (SG vs. SDG) as a between-participants factor. This analysis revealed a significant main effect of session for all conditions (O, F(1,260)=73.160, η^2^=.21; U, F(1,252)=48.889, η^2^=.16; L, F(1,260)=17.852, η^2^=.06, all *ps* < .001) whereby performance significantly improved from training to retest. A significant main effect of group was observed for the O condition (O, F(1,260)=21.360, *p* < .001, η^2^=.06) where performance was faster in the SDG as compared to the SG. No such group effects were observed for the U and L conditions (U, F(1,252)=1.089, η^2^=.004; L, F(1,260)=0.001, η^2^=.000002, all ps > .2). The group by session interaction was not significant in any of the conditions (O, F(1,260)=0.001, η^2^=.000002; U, F(1,252)=0.08, η^2^=.0003; L, F(1,260)=0.075, η^2^=.0003, all *ps* > .7; Figure 2B) suggesting that sleep in the first post-training night did not influence offline changes in performance in any of the conditions.

### Brain activity

#### Finger tapping is associated with motor beta ERD and inter-tapping rest periods with motor-parietal beta ERS

We first aimed to identify finger tapping-ERD patterns at the source level. To do so, we used non- parametric tests (as implemented in Fieldtrip with cluster-based correction in time and space) of the data of all conditions to contrast the post-button-press period (0 to 500ms) and a 500ms resting state period.

This analysis revealed an ERD in the beta range over right motor regions (M1) between 100 and 300 ms after the first correct key press of a chunk (alpha = .05, k = 5549 vertices, area = 899.26 cm^2^, *p* < .001; Figure 3A). In the subsequent analyses, source-localised beta activity related to task practice was examined within this 100-300ms time-window.

**Figure 3.**
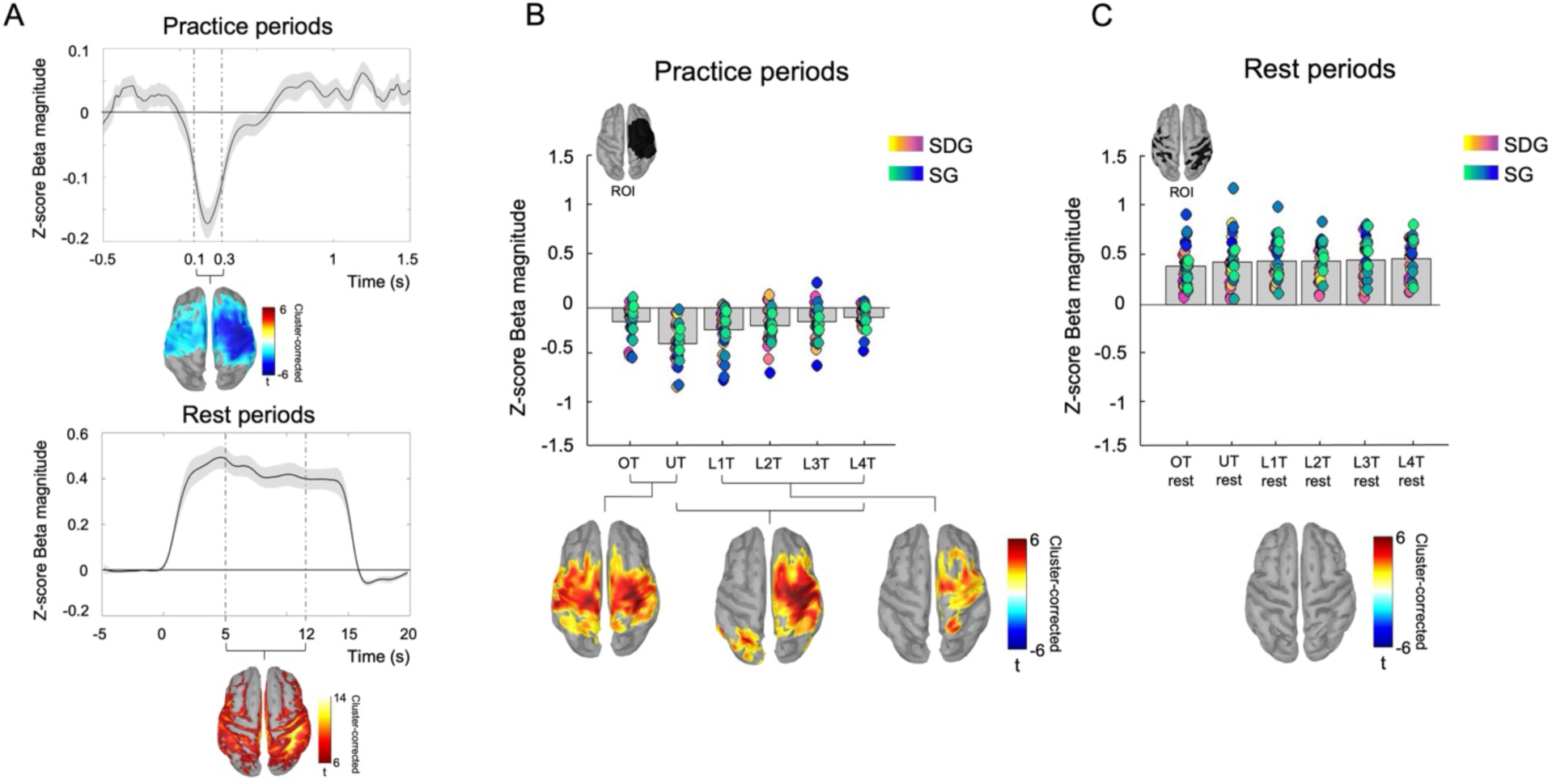
Beta oscillations associated to task practice are modulated by task conditions and learning stage during initial training. **A:** Time series of beta oscillations (18-25 Hz) in motor regions during practice from -500 ms to 1500 ms relative to the first button press of a correct chunk (top panel) and in parietal regions during inter-practice rest period from -5000 ms to 20000 ms relative to the beginning of the rest period (bottom panel). The average of all conditions is represented by the black line, with the shaded grey representing the SEM. Vertical doted lines indicate the significant (cluster corrected in time and space) time periods where ERS/ERD emerged and that were used for the subsequent analyses (100- 300ms for practice periods and 5-12s for inter-practice rest periods). The significant spatial clusters are displayed on MNI-152 cortical mesh provided by SPM12 at a significance level α of .05. These spatial clusters are used for all subsequent analyses to extract beta estimates for illustration purposes. **B:** Bar plot represents beta magnitude (Z-score) in the practice ROI defined in panel A of all participants for each condition during the training session with colored circles indicating individual participants. Source maps represent significant contrasts (cluster corrected in space) performed at the whole brain level, displayed on MNI-152 cortical mesh provided by SPM12 at a significance level α of .05. Results show a progressive decrease of beta desynchronization over contralateral motor regions throughout learning. **C:** Bar plot represents beta magnitude (Z-score) in the inter-practice rest period ROI defined in panel A of all participants for rest periods following each condition during the training session. Colored circles indicate individual participants. The source map illustrates that no contrasts (cluster corrected) were significant (performed at the whole brain level, displayed on MNI-152 cortical mesh provided by SPM12 at a significance level α of .05).

We then tested whether beta ERS patterns could be identified during inter-tapping rest periods. To do so, the data of all inter-practice rest periods (irrespective of the condition it followed) were used to perform a cluster-corrected contrast at the whole brain level between the rest period (2000 to 14000 ms) and the pre-rest baseline period (-4000 to 0ms). This analysis revealed a main effect over parietal regions, characterized by a beta synchronization (ERS) during the rest period, from 2 to 14 seconds (alpha = .05, k = 337 vertices, area = 55.35 cm2, *p* = .002; Figure 3A). In the subsequent analyses, source-localised beta activity during rest periods was examined within a narrower window (5-12s) to avoid the edge effects due to the transition in beta activity between practice and rest periods.

#### Progressive reduction of motor beta ERD as a function of motor sequence practice during initial training

We investigated whether practice-related ERDs were modulated by task condition and learning during initial training. Using cluster-based permutation in the spatial domain, we first contrasted data from the U condition (UT) with data from the O condition (OT) during the period of interest (100 to 300ms relative to the first button press of a correct chunk) identified in Figure 3A. We observed that in bilateral motor regions, beta desynchronization was greater for the more complex and novel UT condition as compared to the more automatized OT condition (left cluster: alpha = .05, k = 1775 vertices, area = 285.41 cm^2^; right cluster: alpha = .05, k = 1467 vertices, area = 236.08 cm^2^; all *p*s = .001, Figure 3B). We then investigated the main effect of learning during training by comparing the electrophysiological responses of the LTs condition versus the UT condition. Cluster-based permutation tests showed that during movement execution, the magnitude of the desynchronization progressively and significantly decreased during practice of the LT sequence as compared to UT in motor regions contralateral to the hand used for the task (e.g., UT vs. L4T, left cluster: alpha = .05, k = 438 vertices, area = 67.29 cm^2^, p < .05; right cluster: alpha = .05, k = 1715 vertices, area = 278.17 cm^2^, *p* = .001, Figure 3B; and see supplementary Figure 2 for detailed comparison between UT and all LT conditions). A similar effect was observed throughout the repetition of the LT sequence (i.e., contrasting electrophysiological data of each LT condition with one another: L1T, L2T, L3T, L4T). Specifically, this analysis revealed that at the end of the training session (L4T), beta desynchronization associated to motor execution during practice was reduced as compared to the beginning of learning in contralateral motor regions (L4T vs. L1T, alpha = .05, k = 919 vertices, area = 150.29 cm^2^, p < .01, Figure 3B). However, there were no significant differences between each L conditions throughout practice (L1T vs. L2T, L2T vs. L3T, L3T vs. L4T; all *ps* > 0.1). As expected, there were no significant differences between groups in any of the contrast reported above (all *ps* > 0.1). Altogether, these results suggest that movement-related beta desynchronization significantly decreased as learning progressed and participants became faster at performing the motor sequence task.

#### ERD modulation is learning-specific in the left M1 but not the right M1

To examine the influence of performance speed on the effects reported above, we compared OT to all LTs, in which different speed was observed. When comparing L1T to OT, we observed a significant difference in bilateral motor areas (OT vs L1T, left cluster: alpha = .05, k = 1276 vertices, area = 208.02 cm^2^, *p* < .01; right cluster: alpha = .05, k = 584 vertices, area = 92.24 cm^2^, *p* < .05). When comparing OT with other LTs, there was no difference in power in the contralateral motor areas (see beta extracted from the practice ROI in Figure 3A), but a significant modulation was observed in the left (ipsilateral) motor cortical areas, such that practice of the L sequence resulted in greater desynchronization than the O sequence (OT vs L2T, alpha = .05, k = 771 vertices, area = 126.42 cm^2^, *p* < .05 ; OT vs L3T, alpha = .05, k = 885 vertices, area = 137.86 cm^2^, *p* < .05 ; OT vs L4T, alpha = .05, k = 841 vertices, area = 135 cm^2^, *p* < .05). The effect on the left M1 was observed consistently irrespective of differences in speed in the different L blocks. These effects did not differ between groups (all *ps* > 0.1). In sum, these modulations are consistent with the effects described above, i.e., greater ERD for the more complex L condition than for the O condition, even when performance speed varies between conditions (Figure 4).

**Figure 4.**
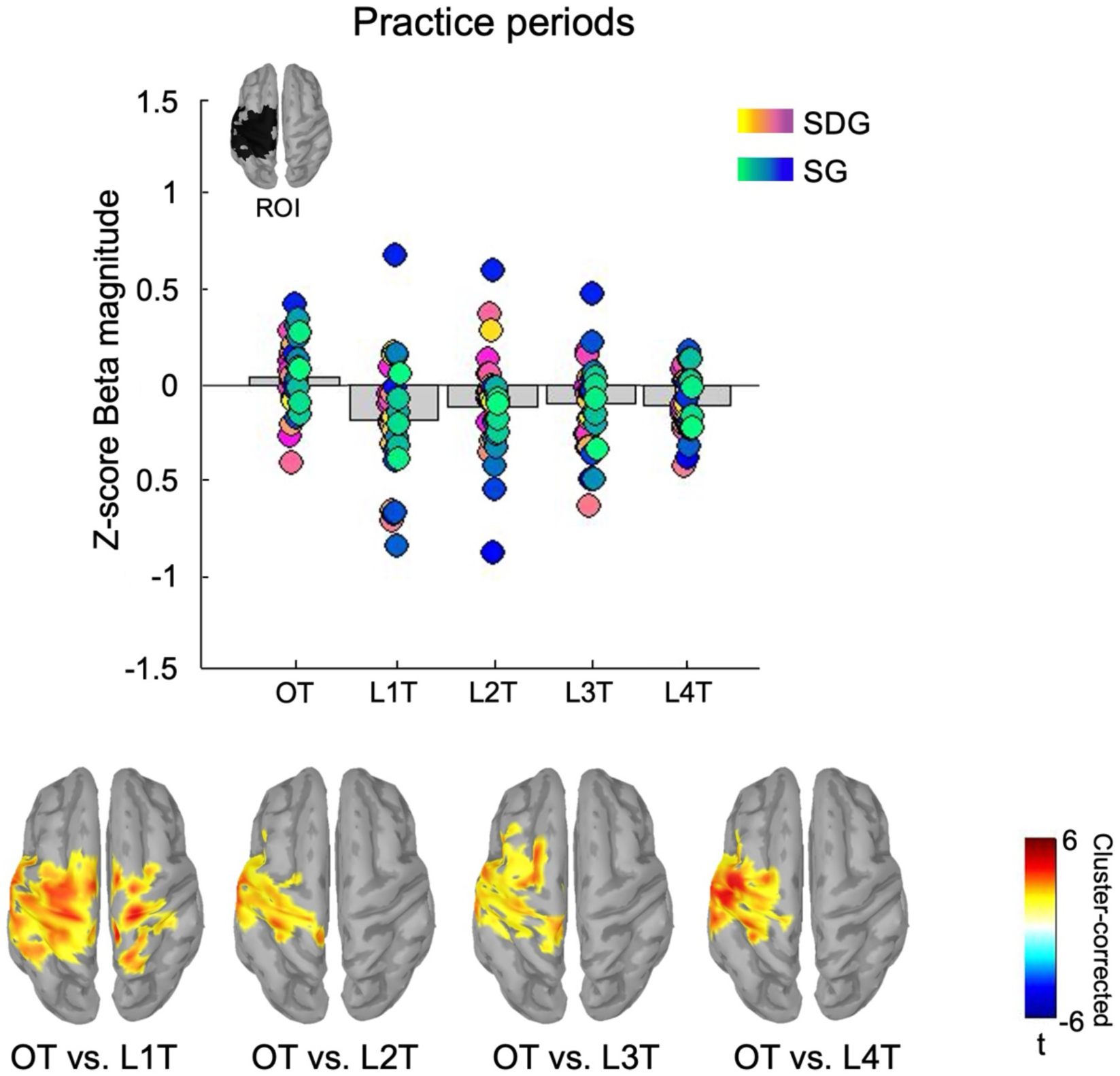
Beta oscillations in left motor regions associated with task practice are modulated by the sequence’s complexity during initial training. Bar plot represents beta magnitude (Z-score) in motor regions (in the left motor cortex, identified with a 15% threshold at the most significant cluster) of all participants for the OT, L1T, L2T, L3T and L4T conditions during the training session, with colored circles indicating individual participants. Source maps represent significant contrasts (cluster corrected) performed at the whole brain level displayed on MNI-152 cortical mesh provided in SPM12 at a significance level α of .05. Results show a greater beta desynchronization in the left M1 during the four blocks of the L sequence as compared to the O sequence during initial training.

#### Beta ERS during rest periods is not modulated by learning or task complexity during initial training

We investigated the modulation of beta activity during inter-practice rest periods following the different conditions with cluster-based permutations tests on the period of interest defined in Figure 3A (lower panel, 5 to 12s relative to the beginning of the rest period). We did not observe any differences in beta magnitude between any of the conditions during rest (*all ps* > .05; see Figure 3C), which suggests that the beta ERS during rest periods interspersed with task practice is not modulated by learning or task complexity during initial training.

#### Beta ERD during practice periods at retest is not modulated by the sleep status of the first post-training night

We then investigated if practice-related changes in oscillatory patterns were modulated during retest depending on the sleep status of the first post-training night. Similar as during initial learning, these analyses showed greater beta ERD in bilateral motor regions for the more complex UR compared to OR sequence in the SG (left cluster: alpha = .05, k = 1232 vertices, area = 196.35 cm^2^, *p* = .002; right cluster: alpha = .05, k = 624 vertices, area = 103.04 cm^2^, *p* < .05), and in contralateral motor regions for the SDG (alpha = .05, k = 1809 vertices, area = 289.47 cm^2^, *p* < .01). However, this effect did not differ between groups (p > .05). Comparisons between LR and UR conditions showed greater beta ERD in bilateral motor regions for UR compared to L1R (but not L2R, L3R and L4R) for the SG (left cluster: alpha = .05, k = 1304 vertices, area = 206.95 cm^2^, *p* = .001; right cluster: alpha = .05, k = 603 vertices, area = 99.30 cm^2^, *p* = .05) while no such difference was observed in the SDG. This effect did not differ between groups (p > .05). Finally, contrasting electrophysiological data of each LR condition with one another (L1R, L2R, L3R, L4R) did not yield any significant effect (Figure 5A). These results suggest that the beta ERD related to motor execution during retest - described during the initial training session to be modulated by learning - was not influenced by the sleep condition of the first post-training night.

**Figure 5.**
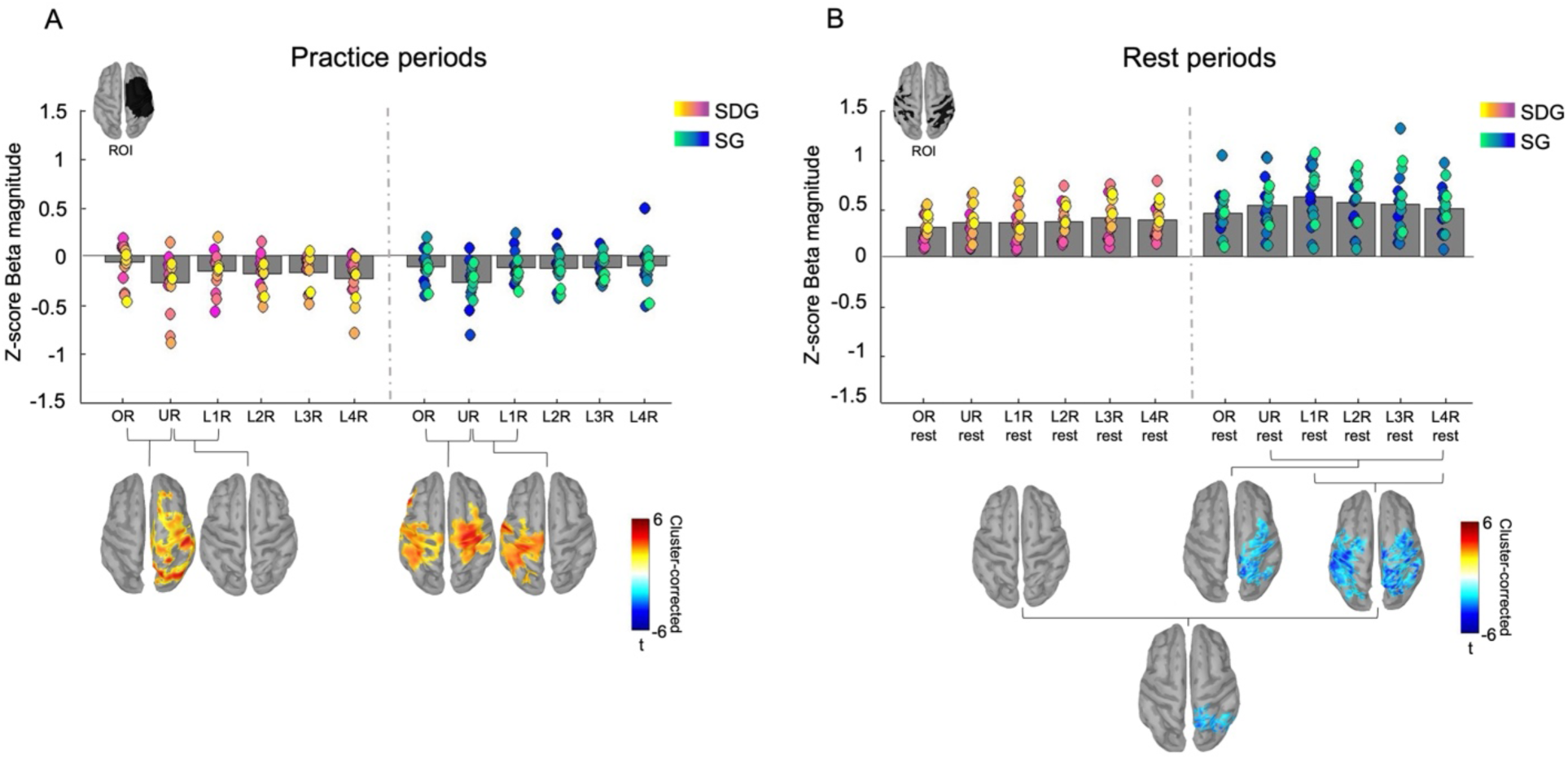
Beta oscillations associated with task and inter-practice rest periods during the retest session in the SG and SDG. **A:** Bar plot represents beta magnitude (Z-score) in motor regions (practice ROI) of all participants for each condition during retest session separated by groups with colored circle indicating individual participants. Source maps represent significant contrasts (cluster corrected) performed at the whole brain level, cortical mesh MNI-152 provided by SPM12 at a significance level α of .05. Task- related ERD modulation did not differ between groups. **B:** Bar plot represents beta magnitude (Z-score) in parietal regions (rest ROI) of all participants for rest periods following each condition during retest session separated by groups with colored circles indicating individual participants. Source maps represent significant contrasts (cluster corrected) performed at the whole brain level, cortical mesh MNI-152 provided by SPM12 at a significance level α of .05, for SG and between the SG and SDG. Results suggest a greater learning-related difference in ERS during rest in the SG as compared to the SDG.

#### Beta ERS during inter-practice rest periods is modulated by the sleep status of the first post-training night

We then examine the effect of sleep on beta ERS during the rest periods in the retest session. We observed a significant difference between beta ERS during the rest periods following the practice of the U condition (URrest) and the last four blocks of the L condition (L4Rrest). Specifically, results show a decreased beta ERS in L4Rrest as compared to URrest for the SG (alpha = .05, k = 576 vertices, area = 96.10 cm^2^, *p* < .05). Such effect was not observed in the SDG, but the group contrast was not significant (*p* > .05). Moreover, we observed a significant difference between the last four blocks of the L condition (L4Rrest) and the first four blocks (L1Rrest), where the beta ERS decreases at the end of learning (L4Rrest) as compared to the beginning (L1Rrest) in motor and parietal regions. This result was observed only for the SG (left cluster: alpha = .05, k = 738 vertices, area = 123.57 cm^2^, *p* = .01; right cluster: alpha = .05, k = 892 vertices, area = 146.22 cm^2^, *p* < .01) but not in the SDG (*p* > .05). This effect was greater for the SG as compared to SDG in contralateral parietal regions (alpha = .05, k = 406 vertices, area = 63.60 cm^2^, *p* = .05; Figure 5B). Data inspection suggests that these effects were driven by heightened ERS after L1R in the sleep group. Altogether, these results suggest that sleep (as compared to sleep deprivation) after initial learning resulted in a greater difference in beta ERS during inter-practice rest periods following the learned sequence early during the 72h retest session (L1R).

#### Offline changes in beta ERD during practice are not modulated by a night of sleep

To assess the between-session changes in oscillatory activity associated to practice, we contrasted data of the retest session with data of the training session for the L condition (L1R vs L4T; Figure 6A), the O condition and the U condition (OR vs OT, and UR vs UT; Supplementary Figure 3) for each group separately. During practice, no significant difference emerged between sessions in any of the groups and any of all conditions (*p* > .1).

**Figure 6.**
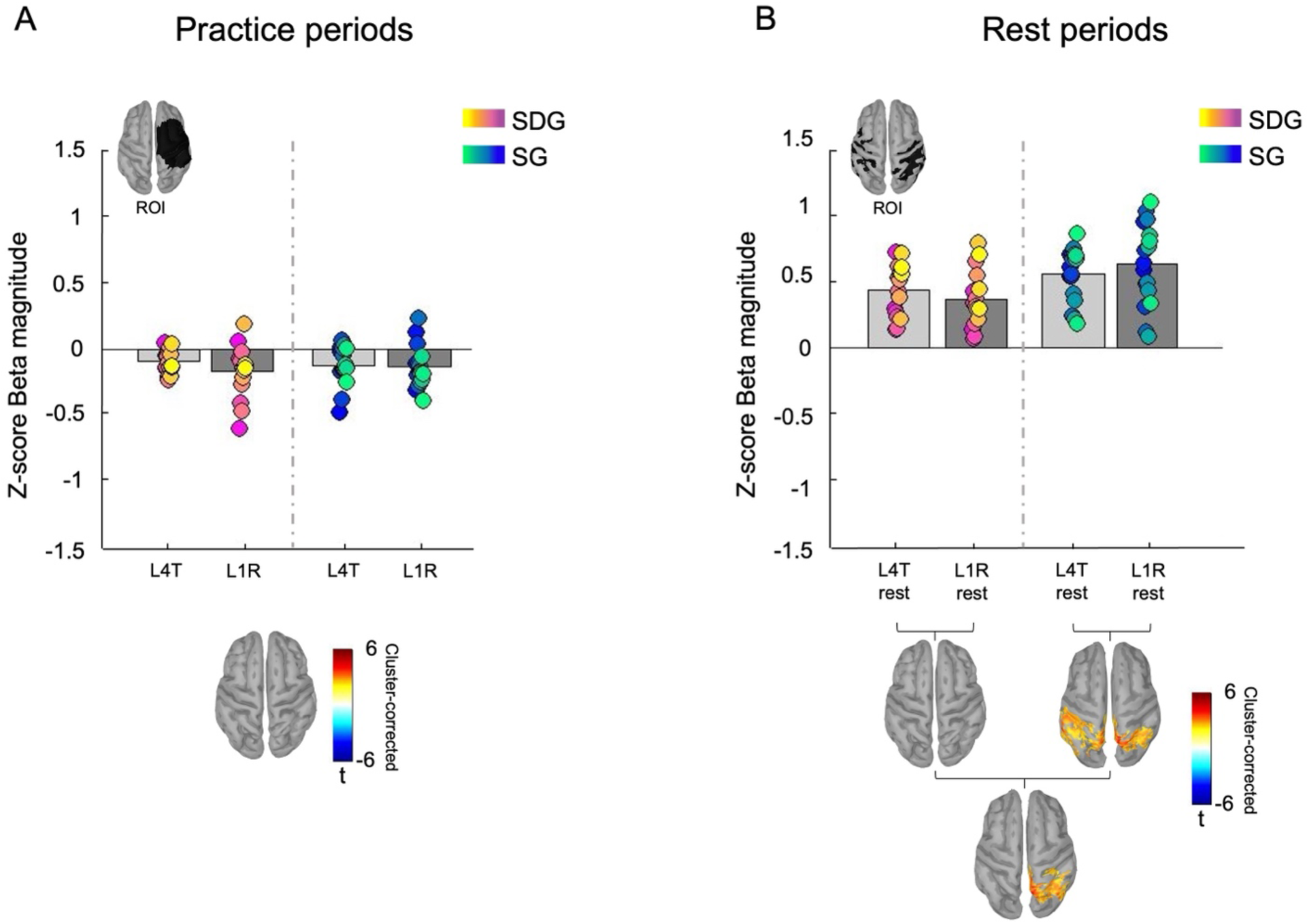
Between-session changes in beta oscillations during practice and inter-practice rest periods in the SG and the SDG. **A:** Bar plot represents beta magnitude (Z-score) in motor regions (practice ROI) of all participants for L4T and L1R conditions during practice separated by groups with colored circles indicating individual participants. Source maps represent significant contrasts (cluster corrected) performed at the whole brain level cortical mesh MNI-152 provided by SPM12 at a significance level α of .05. No significant cluster emerged from whole brain contrasts between conditions. **B:** Bar plot represents beta magnitude (Z-score) in parietal regions (rest ROI) of all participants for the rest periods following L4T and L1R conditions separated by groups with colored circles indicating individual participants. Source maps represent significant contrasts (cluster corrected) performed at the whole brain level cortical mesh MNI-152 provided by SPM12 at a significance level α of .05. Results suggest a greater beta synchronization during the rest period of the L1R as compared to the rest period of the L4T, and only for SG.

#### Offline changes in beta ERS during inter-practice rest periods are modulated by a night of sleep

To evaluate the between-session changes in oscillatory activity during inter-practice rest periods, we contrasted rest data of the retest session with rest data of the training session for the L condition (L1Rrest vs L4Trest; Figure 6A), the O condition and the U condition (ORrest vs OTrest, and URrest vs UTrest; Supplementary Figure 3) for each group separately. No significant session effect was observed between O conditions and U conditions (p > .1; Supplementary Figure 3). Interestingly, a significant difference between L1Rrest and L4Trest was observed for the SG, such that the beta ERS was greater at retest as compared to training (left cluster: alpha = .05, k = 947 vertices, area = 148.09 cm^2^; right cluster: alpha= .05, k = 586 vertices, area = 90.15 cm^2^; *all ps* < .05) in bilateral parietal regions. No such effect was observed between sessions for the SDG (*p* > .05). Finally, this effect was significantly different between groups in contralateral parietal regions (greater ERS at retest vs. training for SG as compared to SDG; alpha = .05, k = 656 vertices, area = 100 cm^2^, *p* < .05, Figure 6B).

Finally, we performed Pearson’s correlations to estimate the relationship between learning- and sleep-related modulations in beta oscillatory activity during practice (and inter-practice rest periods) at the source level and offline changes in performance (accuracy and speed). No significant correlation was observed in any of the two groups for any conditions (FDR corrected).

## Discussion

This study investigates the brain oscillatory signatures of motor sequence learning and the impact of post-learning sleep (SG) - as compared to sleep deprivation (SDG) - on these signatures in young healthy participants using MEG. Results indicate that initial learning was paralleled by a progressive decrease in event-related desynchronization (ERD) in the beta band (18–25 Hz) over motor areas during task practice while event-related synchronization (ERS) remained stable in the same band throughout learning. While motor performance improved over the consolidation period regardless of the sleep condition, beta ERS during the short inter-practice rest periods following the learned sequence were specifically enhanced over bilateral parietal regions in the SG as compared to the SGD group early during retest. Altogether, these results suggest that sleep after motor learning modulates the amplitude of brain synchronization over parietal regions following the practice of learned movements.

### No evidence for a beneficial effect of sleep as compared to sleep deprivation on motor performance

Participants showed significant offline gains in performance on the learned motor sequence across the consolidation interval, but contrary to our expectations, this was observed regardless of whether they had a full night of sleep or were totally sleep deprived after initial learning. While this is in line with a prior study using the same motor task reporting no significant effect of sleep deprivation on motor performance (Albouy et al. 2013b), this observation stands in contrast with earlier studies showing sleep-dependent motor memory consolidation (see King et al. 2017 for a review).

One plausible explanation for the lack of differences between groups is the affordance of the two recovery nights to the SDG to minimize the effect of fatigue on performance at retest, which may have allowed for delayed performance improvements and thus reduced the effect of the sleep deprivation on performance (Albouy et al. 2013b; Walker et al. 2003). It is also possible that our specific retest design may have masked the potential beneficial effect of sleep on performance. Specifically, during the retest session, participants first practiced the overlearned and untrained sequences before being retested on the learned sequence. Potential pro-active interference of these conditions on the subsequently performed learned sequence could have reduced our ability to detect a more subtle beneficial effect of sleep on performance. Alternatively, the absence of a sleep-dependent effect on motor performance concurs with recent models challenging the view that sleep (as compared to wakefulness) plays a beneficial role in the consolidation of motor memories (Rickard et al. 2022). The current design can’t tease apart these different alternatives.

### Initial motor sequence learning - but not post-learning sleep - modulates the amplitude of beta ERD over the left M1

As expected, MEG data analyses showed that the execution of a movement - irrespective of the condition – was associated with an ERD over motor areas (Neuper et al. 2006; Pfurtscheller and Lopes da Silva 1999). Interestingly, this marker of motor task practice was modulated by the task condition. Specifically, during practice of the unlearned condition, beta ERD was greater in bilateral motor regions as compared to the simpler overlearned condition. This result suggests that beta ERD is modulated by the complexity of the sequence. This is in line with prior findings suggesting that the practice of finger-tapping sequences elicit greater M1 recruitment (Gerloff et al. 1998) and stronger ERD (Manganotti et al. 1998; Hummel et al. 2003) with increasing sequence complexity. These effects have been attributed to the greater preparation and attentional processes necessary for fast and accurate execution of complex – as compared to simple – sequences of movement (Hummel et al. 2003; Manganotti et al. 1998).

Interestingly, a progressive decrease in beta ERD amplitude was observed over bilateral motor regions as a function of practice of the learned sequence. In order to test the extent to which this effect is related to differences in performance speed between early and late training (i.e., L1T vs. L4T) or to differences in learning stage, we contrasted oscillatory brain activity between learning condition and the simple overlearned sequence (LTs vs. O) that presented different performance speed. The results indicate a difference in ERD amplitude over the left M1 - but not the right M1 – irrespective of performance speed (Figure 3). These findings suggest that the ERD modulations observed between early vs. late training reported over the left M1 might be related to learning *per se* while those observed on the right M1 might be related to differences in performance speed between early and late training blocks. These findings are in line with previous literature suggesting that left hemisphere, but not right hemisphere stroke is associated with deficits in the encoding and generation of sequential movements (Harrington and Haaland 1991; Kimura 1977). Additionally, non-invasive brain stimulation studies showed that left M1 stimulation benefits motor learning (performed with the right hand) more than sham or right M1 stimulation (Schambra et al. 2011). Overall, our results therefore suggest that the amplitude of beta ERD over the left M1 is specifically modulated by the motor sequence learning process.

As our results suggest that beta ERD over the left M1 is an oscillatory marker of the motor learning process, we examined whether this signature was modulated by post-learning sleep. Our results did not indicate any effect of the sleep condition on beta ERD amplitude at retest. These results are consistent with recent models suggesting that M1 does not play a critical role in sleep-dependent motor sequence memory consolidation (King et al. 2017). Interestingly, there was also no evidence for a modulation of beta ERD as a function of (re)learning during retest or for changes in ERD amplitude between training and retest. These results suggest that the modulation of beta ERD observed over the left M1 during training was specific to the initial – early – learning phase. These results are in line with previous studies proposing that M1 supports performance during the early phases of learning but not in the delayed stages of motor skill consolidation (Robertson et al. 2005; Hotermans et al. 2008). Altogether, these findings concur with views that beta ERD are neurophysiological markers of early cortical reorganization associated with motor learning (Pollok et al. 2014).

### Inter-practice rest ERS is not modulated by initial training but by post-learning sleep

Our MEG data analyses revealed the presence of beta ERS over bilateral parietal and motor regions that were sustained during the entire duration of the inter-practice rest periods during initial learning. These results are in line with previous studies reporting an increase in the magnitude of beta oscillatory activity during post-movement rest periods, as compared to during movement time, over the brain regions involved during task practice (Jurkiewicz et al. 2006; Pfurtscheller and Lopes da Silva 1999; Neuper et al. 2006). While beta ERS has usually been associated with active motor inhibition during rest periods following movements (Heinrichs-Graham et al. 2017), more recent research has proposed that oscillatory brain activity during inter-practice rest periods might reflect an active memory consolidation process (Buch et al. 2021; Bönstrup et al. 2019). Specifically, recent studies found that contralateral frontoparietal beta oscillatory activity (Bönstrup et al. 2019) and task-related neural replay in sensorimotor, entorhinal and hippocampal regions (Buch et al. 2021) are related with improvement in performance occurring over inter-practice rest periods. These improvements have been reported to be particularly pronounced early during training (Buch et al. 2021; Bönstrup et al. 2019), so one could expect the corresponding neural signatures to be modulated by practice / learning stage. However, such modulations of brain activity throughout learning were not examined in these earlier studies and the current results do not support this view as the amplitude of the beta ERS observed in our dataset remained consistent throughout initial learning.

While learning and conditions did not modulate beta ERS during initial training, post-learning sleep precisely influenced this oscillatory marker. Specifically, we observed a sleep-dependent increase in the amplitude of beta ERS during inter-tapping rest periods during early retest as compared to the end of training over parietal regions. This sleep-dependent boost in beta ERS over parietal regions was observed when participants were firstly reintroduced to the previously learned sequence (L1R) after sleep and this effect was followed by a linear decrease of beta ERS amplitude throughout the retest on the L condition (see L1R vs. L4R contrast). No such effects were observed in the sleep-deprived group. These results suggest that ERS might be an oscillatory marker of sleep-dependent consolidation that appears to be modulated at later stages of learning. This stands in contrast with earlier reports suggesting that oscillatory brain activity during inter-practice rest reflects early and fast consolidation processes (Buch et al. 2021; Bönstrup et al. 2019). Here, we suggest that the amplitude of the post-movement beta rebound over parietal regions is influenced by sleep during the slow consolidation process. While task-related brain activity in parietal brain regions has been described to decrease as a function of learning (Doyon et al. 2009) and after a night of sleep (Walker et al. 2005), there is evidence to suggest that the parietal cortex support motor memory consolidation during sleep. Specifically, an increase in slow wave activity was reported over parietal regions during sleep following motor learning (Huber et al. 2004). This modulation in parietal oscillatory activity was linked to performance improvement the next day (note though that this study employed an adaptation, but not a MSL, task). In a more recent review (King et al. 2017), it was suggested that spatial representations of motor sequences encoded in hippocampal-parietal networks during learning might be reactivated during post-learning sleep and strengthen the motor memory consolidation process. In the same vein, but on a different timescale, Buch and collaborators (2021) showed evidence for reactivations of task-related patterns in hippocampo-parietal networks during short inter-practice offline rest episodes during initial learning. Taken together with this earlier work, our findings suggest that newly encoded motor memory traces might have been reactivated in (hippocampo-)parietal networks during post-learning sleep and that this process might be reflected by a boost in parietal ERS during the short offline rest episodes following the practice of these consolidation motor sequences. However, this remains speculative and further investigation into the oscillatory processes supporting motor memory consolidation during post-learning sleep is warranted.

## Conclusion

Our results show that initial motor sequence learning was paralleled by a progressive decrease in event-related desynchronization in the beta band over motor areas during task practice, but these neural signatures of motor learning were not modulated by post-learning sleep. In contrast, event-related synchronization during the short inter-practice rest periods following the learned sequence was specifically enhanced after sleep over bilateral parietal regions. Altogether, these results suggest that sleep enhances the synchronisation of parietal brain oscillations following the practice of learned movements, even in the absence of overt behavioral changes.

## Acknowledgments

This work was supported by a grant from Fonds de Recherche du Québec – Santé and Brain Canada Future Leaders to P.A., a grant from Réseau de Bio-Imagerie du Québec to A.S. and P.A., a grant from Centre de Recherche du Québec Audace to P.A., and NSERC Discovery grants to P.A. This study was also supported by the C.N.R.S. (Centre National de la Recherche Scientifique), the I.N.S.E.R.M. (Institut National de la Santé Et de la Recherche Médicale), the University of Lyon and the Région Rhône-Alpes.

## Author contributions

Marie-Frédérique Bernier (Data curation, Formal analysis, Investigation, Methodology, Visualization, Writing – original draft, Writing – review & editing), Roxane Hoyer (Data curation, Formal analysis, Methodology, Writing – review & editing), Françoise Lecaignard (Data curation, Formal analysis, Investigation, Methodology), Alain Nicolas (Methodology, Project administration, Resources), Olivier Bertrand (Conceptualization, Data curation, Funding acquisition, Methodology, Resources, Writing – review & editing), Philippe Albouy (Conceptualization, Data curation, Formal analysis, Funding acquisition, Methodology, Project administration, Resources, Supervision, Validation, Writing – review & editing), and Geneviève Albouy (Conceptualization, Data curation, Formal analysis, Methodology, Project administration, Resources, Supervision, Validation, Writing – review & editing).

## Data availability

Source data used to produce the figures and tables will be made available upon publication.

## References

Albouy G, Fogel S, King BR, Laventure S, Benali H, Karni A, Carrier J, Robertson EM, Doyon J. 2015. Maintaining vs. enhancing motor sequence memories: respective roles of striatal and hippocampal systems. NeuroImage. 108:423–434. doi:10.1016/j.neuroimage.2014.12.049

Albouy G, King BR, Maquet P, Doyon J. 2013a. Hippocampus and striatum: dynamics and interaction during acquisition and sleep-related motor sequence memory consolidation. Hippocampus. 23(11):985–1004. doi:10.1002/hipo.22183

Albouy G, Sterpenich V, Vandewalle G, Darsaud A, Gais S, Rauchs G, Desseilles M, Boly M, Dang-Vu T, Balteau E, Degueldre C, Phillips C, Luxen A, Maquet P. 2013b. Interaction between hippocampal and striatal systems predicts subsequent consolidation of motor sequence memory. PLoS One. 8(3):e59490. doi:10.1371/journal.pone.0059490

Baillet S, Mosher JC, Leahy RM. 2001. Electromagnetic brain mapping. IEEE Signal Process Mag. 18(6):14–30. doi:10.1109/79.962275

Beck AT. 1961. An inventory for measuring depression. Arch Gen Psychiatry. 4(6):561. doi:10.1001/archpsyc.1961.01710120031004

Beck AT, Epstein N, Brown G, Steer RA. 1988. An inventory for measuring clinical anxiety: psychometric properties. J Consult Clin Psychol. 56(6):893–897. doi:10.1037/0022-006X.56.6.893

Bönstrup M, Iturrate I, Thompson R, Cruciani G, Censor N, Cohen LG. 2019. A rapid form of offline consolidation in skill learning. Curr Biol. 29(8):1346–1351.e4. doi:10.1016/j.cub.2019.02.049

Buch ER, Claudino L, Quentin R, Bönstrup M, Cohen LG. 2021. Consolidation of human skill linked to waking hippocampo-neocortical replay. Cell Rep. 35(10):109193. doi:10.1016/j.celrep.2021.109193

Doyon J, Bellec P, Amsel R, Penhune V, Monchi O, Carrier J, Lehéricy S, Benali H. 2009. Contributions of the basal ganglia and functionally related brain structures to motor learning. Behav Brain Res. 199(1):61–75. doi:10.1016/j.bbr.2008.11.012

Dyck S, Klaes C. 2024. Training-related changes in neural beta oscillations associated with implicit and explicit motor sequence learning. Sci Rep. 14(1):6781. doi:10.1038/s41598-024-57285-7

Ellis BW, Johns MW, Lancaster R, Raptopoulos P, Angelopoulos N, Priest RG. 1981. The St. Mary’s Hospital Sleep Questionnaire: a study of reliability. Sleep. 4(1):93–97. doi:10.1093/sleep/4.1.93

Formaggio E, Storti SF, Avesani M, Cerini R, Milanese F, Gasparini A, Acler M, Pozzi Mucelli R, Fiaschi A, Manganotti P. 2008. EEG and fMRI coregistration to investigate the cortical oscillatory activities during finger movement. Brain Topogr. 21(2):100–111. doi:10.1007/s10548-008-0058-1

Gerloff C, Corwell B, Chen R, Hallett M, Cohen LG. 1998. The role of the human motor cortex in the control of complex and simple finger movement sequences. Brain. 121(9):1695–1709. doi:10.1093/brain/121.9.1695

Harrington DL, Haaland KY. 1991. Hemispheric specialization for motor sequencing: abnormalities in levels of programming. Neuropsychologia. 29(2):147–163. doi:10.1016/0028-3932(91)90017-3

Heinrichs-Graham E, Kurz MJ, Gehringer JE, Wilson TW. 2017. The functional role of post-movement beta oscillations in motor termination. Brain Struct Funct. 222(7):3075–3086. doi:10.1007/s00429-017-1387-1

Horne JA, Ostberg O. 1976. A self-assessment questionnaire to determine morningness-eveningness in human circadian rhythms. Int J Chronobiol. 4(2):97–110.

Hotermans C, Peigneux P, De Noordhout AM, Moonen G, Maquet P. 2008. Repetitive transcranial magnetic stimulation over the primary motor cortex disrupts early boost but not delayed gains in performance in motor sequence learning. Eur J Neurosci. 28(6):1216–1221. doi:10.1111/j.1460-9568.2008.06421.x

Huang MX, Mosher JC, Leahy RM. 1999. A sensor-weighted overlapping-sphere head model and exhaustive head model comparison for MEG. Phys Med Biol. 44(2):423–440. doi:10.1088/0031-9155/44/2/010

Huber R, Felice Ghilardi M, Massimini M, Tononi G. 2004. Local sleep and learning. Nature. 430(6995):78–81. doi:10.1038/nature02663

Hummel F, Kirsammer R, Gerloff C. 2003. Ipsilateral cortical activation during finger sequences of increasing complexity: representation of movement difficulty or memory load? Clin Neurophysiol. 114(4):605–613. doi:10.1016/S1388-2457(02)00417-0

Johns MW. 1991. A new method for measuring daytime sleepiness: the Epworth Sleepiness Scale. Sleep. 14(6):540–545. doi:10.1093/sleep/14.6.540

Jurkiewicz MT, Gaetz WC, Bostan AC, Cheyne D. 2006. Post-movement beta rebound is generated in motor cortex: evidence from neuromagnetic recordings. NeuroImage. 32(3):1281–1289. doi:10.1016/j.neuroimage.2006.06.005

Karni A, Meyer G, Jezzard P, Adams MM, Turner R, Ungerleider LG. 1995. Functional MRI evidence for adult motor cortex plasticity during motor skill learning. Nature. 377(6545):155–158. doi:10.1038/377155a0

Kimura D. 1977. Acquisition of a motor skill after left-hemisphere damage. Brain. 100:527–542.

King BR, Hoedlmoser K, Hirschauer F, Dolfen N, Albouy G. 2017. Sleeping on the motor engram: the multifaceted nature of sleep-related motor memory consolidation. Neurosci Biobehav Rev. 80:1–22. doi:10.1016/j.neubiorev.2017.04.026

Korman M, Doyon J, Doljansky J, Carrier J, Dagan Y, Karni A. 2007. Daytime sleep condenses the time course of motor memory consolidation. Nat Neurosci. 10(9):1206–1213. doi:10.1038/nn1959

Korman M, Raz N, Flash T, Karni A. 2003. Multiple shifts in the representation of a motor sequence during the acquisition of skilled performance. Proc Natl Acad Sci USA. 100(21):12492–12497. doi:10.1073/pnas.2035019100

Manganotti P, Gerloff C, Toro C, Katsuta H, Sadato N, Zhuang P, Leocani L, Hallett M. 1998. Task-related coherence and task-related spectral power changes during sequential finger movements. Electroencephalogr Clin Neurophysiol. 109(1):50–62. doi:10.1016/S0924-980X(97)00074-X

Neuper C, Wörtz M, Pfurtscheller G. 2006. ERD/ERS patterns reflecting sensorimotor activation and deactivation. In: Progress in Brain Research. Vol. 159. p. 211–222. doi:10.1016/S0079-6123(06)59014-4

Oldfield RC. 1971. The assessment and analysis of handedness: the Edinburgh inventory. Neuropsychologia. 9(1):97–113. doi:10.1016/0028-3932(71)90067-4

Oostenveld R, Fries P, Maris E, Schoffelen JM. 2011. FieldTrip: open source software for advanced analysis of MEG, EEG, and invasive electrophysiological data. Comput Intell Neurosci. 2011:1–9. doi:10.1155/2011/156869

Penhune VB, Steele CJ. 2012. Parallel contributions of cerebellar, striatal and M1 mechanisms to motor sequence learning. Behav Brain Res. 226(2):579–591. doi:10.1016/j.bbr.2011.09.044

Pfurtscheller G. 2001. Functional brain imaging based on ERD/ERS. Vision Res. 41(10–11):1257–1260. doi:10.1016/S0042-6989(00)00235-2

Pfurtscheller G, Lopes da Silva FH. 1999. Event-related EEG/MEG synchronization and desynchronization: basic principles. Clin Neurophysiol. 110(11):1842–1857. doi:10.1016/S1388-2457(99)00141-8

Pineda JA. 2005. The functional significance of mu rhythms: translating “seeing” and “hearing” into “doing.” Brain Res Rev. 50(1):57–68. doi:10.1016/j.brainresrev.2005.04.005

Pollok B, Latz D, Krause V, Butz M, Schnitzler A. 2014. Changes of motor-cortical oscillations associated with motor learning. Neuroscience. 275:47–53. doi:10.1016/j.neuroscience.2014.06.008

Rickard TC, Pan SC, Gupta MW. 2022. Severe publication bias contributes to illusory sleep consolidation in the motor sequence learning literature. J Exp Psychol Learn Mem Cogn. 48(12):1787–1796. doi:10.1037/xlm0001090

Robertson EM, Pascual-Leone A, Miall RC. 2004. Current concepts in procedural consolidation. Nat Rev Neurosci. 5(7):576–582. doi:10.1038/nrn1426

Robertson EM, Press DZ, Pascual-Leone A. 2005. Off-line learning and the primary motor cortex. J Neurosci. 25(27):6372–6378. doi:10.1523/JNEUROSCI.1851-05.2005

Schambra HM, Abe M, Luckenbaugh DA, Reis J, Krakauer JW, Cohen LG. 2011. Probing for hemispheric specialization for motor skill learning: a transcranial direct current stimulation study. J Neurophysiol. 106(2):652–661. doi:10.1152/jn.00210.2011

Tadel F, Baillet S, Mosher JC, Pantazis D, Leahy RM. 2011. Brainstorm: a user-friendly application for MEG/EEG analysis. Comput Intell Neurosci. 2011:1–13. doi:10.1155/2011/879716

Walker MP, Brakefield T, Allan Hobson J, Stickgold R. 2003. Dissociable stages of human memory consolidation and reconsolidation. Nature. 425(6958):616–620. doi:10.1038/nature01930

Walker MP, Stickgold R, Alsop D, Gaab N, Schlaug G. 2005. Sleep-dependent motor memory plasticity in the human brain. Neuroscience. 133(4):911–917. doi:10.1016/j.neuroscience.2005.04.007

