## Supplementary figure 1 for "Sleep after Motor Sequence Learning Enhances Post-Movement Parietal Beta Synchronization"

### Supplements

#### *Sleep prior to and during the experiment*

*Duration and quality of sleep*

Analyses of the Pittsburgh Sleep Quality Index (Buysse et al. 1989) showed that the average sleep duration did not differ between groups during the month preceding the study (SG, 7h54 ± 1h10 min; SDG, 7h34 ± 1h13 min; unpaired test-t, $t(33) = -0.91$, $p = 0.36$). Sleep quality [from poor (1) to good (5)] and duration of the night preceding the training and retest sessions were defined in a subjective way using the St. Mary's Hospital sleep questionnaire (Ellis et al. 1981). The sleep duration average and the subjective sleep quality median did not differ between groups during the night before the training session (Duration: SG, 7 h 46 ± 0 h 58 min; SDG 7 h 47 min ± 0 h 57 min; unpaired t-test, $t(33) = -0.04$, $p = 0.96$; Quality: SG, 4; SDG, 4; unpaired t-test, $t(33) = 0.03$, $p = 0.97$) or during the night before the retest session (Duration: SG, 7 h 37 min ± 1 h 22 min; SDG, 8 h 15 min ± 1h 21 min; unpaired t-test, $t(33) = -1.35$, $p = 0.18$; Quality: SG, 4; SDG, 4; unpaired t-test, $t(33) = -0.81$, p 0.41). Altogether, these results indicate that sleep duration and quality did not differ between groups prior to the experimental sessions.

*Actigraphic data*

Actigraphic data were collected by wrist actigraphy (Cambridge Neuroscience, Cambridge, UK) for 6 days (3 days before and after training session). Actigraphic data on the six nights showed significant main effects of group ($F(1,28)=48.25$, $p < 0.001$) and night ($F(5,140)=65.51$, $p < 0.001$) as well as a group by night interaction ($F(5,140)=65.87$, $p < 0.001$). The activity during the first 3 nights did not differ between groups (all $p > 0.12$). As expected, the activity was larger in the SDG than in the SG group during the deprivation night (SG = 23.29 ± 17.69 units, SDG = 263.5 ± 107.73 units, $F(1,28)=77.53$, $p < 0.001$). During the first recovery night, activity in the SDG tended to be lower than in the SG, suggesting a rebound of sleep after the sleep deprivation (SG = 28.73 ± 24.39 units, SDG = 16.39 ± 11.10 units, $F(1,28)=3.02$, $p = 0.09$). This effect was no longer observed on the second recovery night, which preceded the retest session (SG =

23.98 ± 12.06 units, SDG = 29.55 ± 19.67 units, F(1,28)=0.89, p = 0.35), suggesting that two nights were sufficient to recover from the effects of sleep deprivation.

Actigraphic data during daytime (five days) showed no significant main effects of group (F(1,28)=1.74, p = 0.19) and no group-by-day interaction (F(4,112)=0.79, p = 0.53). The activity during the day following the sleep deprivation did not differ between groups (SG = 289.5 ± 87.56 units, SDG = 307.84 ± 90.56 units, F(1,28)=0.11, p = 0.74), suggesting that, as instructed, sleep-deprived participants maintained a day-like schedule the day after the sleep deprivation.

*Accuracy*

The repeated-measures ANOVAs performed on performance accuracy (i.e., the proportion of correct chunk) during training using block as a within-subject factor and group as a between-subject factor showed no significant main effect of block for all conditions (O, F(3,124)=0.097, $\eta^2$=.002; N, F(3,124)=2.646, $\eta^2$=.06; L, F(15,496)=1.096, $\eta^2$=.03, all *ps* > .05). There was no significant effect of group or group by block interaction in any of the conditions (group effect: O, F(1,124)=3.220, $\eta^2$=.02; U, F(1,124)=0.122, $\eta^2$=.0009; L, F(1,496)=0.585, $\eta^2$=.001, all *ps* > .05; group by block interaction: O, F(3,124)=0.208, $\eta^2$=.005; U, F(3,124)=1.111, $\eta^2$=.02; L, F(15,496)=0.797, $\eta^2$=.02, all *ps* > .3), indicating that accuracy remained high (99.11% ± 4.13) and stable in both groups during training for all conditions (see supplementary figure 1A).

The same ANOVAs performed on the retest data showed no significant main effect of block in any of the conditions (O, F(3,124)=0.038, $\eta^2$=.0009; U, F(3,120)=1.652, $\eta^2$=.04; L, F(15,496)=0.562, $\eta^2$=.02, all *ps* > .1). There was no significant main effect of group in O and U conditions (O, F(1,124)=0.582, $\eta^2$=.005; U, F(1,120)=0.175, $\eta^2$=.001, all *ps* > .4). In contrast, there was a significant main effect of group during retest in the L condition (L, F(1,496)=9.855, $\eta^2$=.02, *p* < .01) whereby participants in the SDG group were more accurate than those in the SG group. However, there was no group by block interaction in any of the conditions (O, F(3,124)=0.278, $\eta^2$=.007; U, F(3,120)=0.978, $\eta^2$=.02; L, F(15,496)=0.327, $\eta^2$=.009, all *ps* >

.4). These results indicate that the accuracy stayed high (99.11% ± 5.28) and constant throughout practice for all conditions (see supplementary figure 1B).

The ANOVA using session (training vs. retest) and blocks (4) as within-subject factors and group as between subject factor did not reveal any significant effect of group for both the O and U conditions (O, $F(1,260)=0.164$, $\eta^2=.0006$; U, $F(1,256)=0.001$, $\eta^2=.0000002$, all *ps* > .6) but a significant main effect of group was observed on the L condition (L, $F(1,260)=5.349$, $\eta^2=.02$, $p < .05$) where accuracy was overall greater in the SDG as compared to the SG. Moreover, for all conditions, there was no significant effect of session (O, $F(1,260)=3.564$, $\eta^2=.01$; U, $F(1,256)=3.363$, $\eta^2=.01$; L, $F(1,260)=0.114$, $\eta^2=.0004$, all *ps* > .06) and no significant group by session interaction (O, $F(1,260)=2.832$, $\eta^2=.01$; U, $F(1,256)=0.277$, $\eta^2=.001$; L, $F(1,260)= 1.097$, $\eta^2=.004$, all *ps* > .09). These results indicate that accuracy did not significantly differ between sessions or between groups (see supplementary figure 1B).

**Supplementary figures**

*Supplementary figure 1*

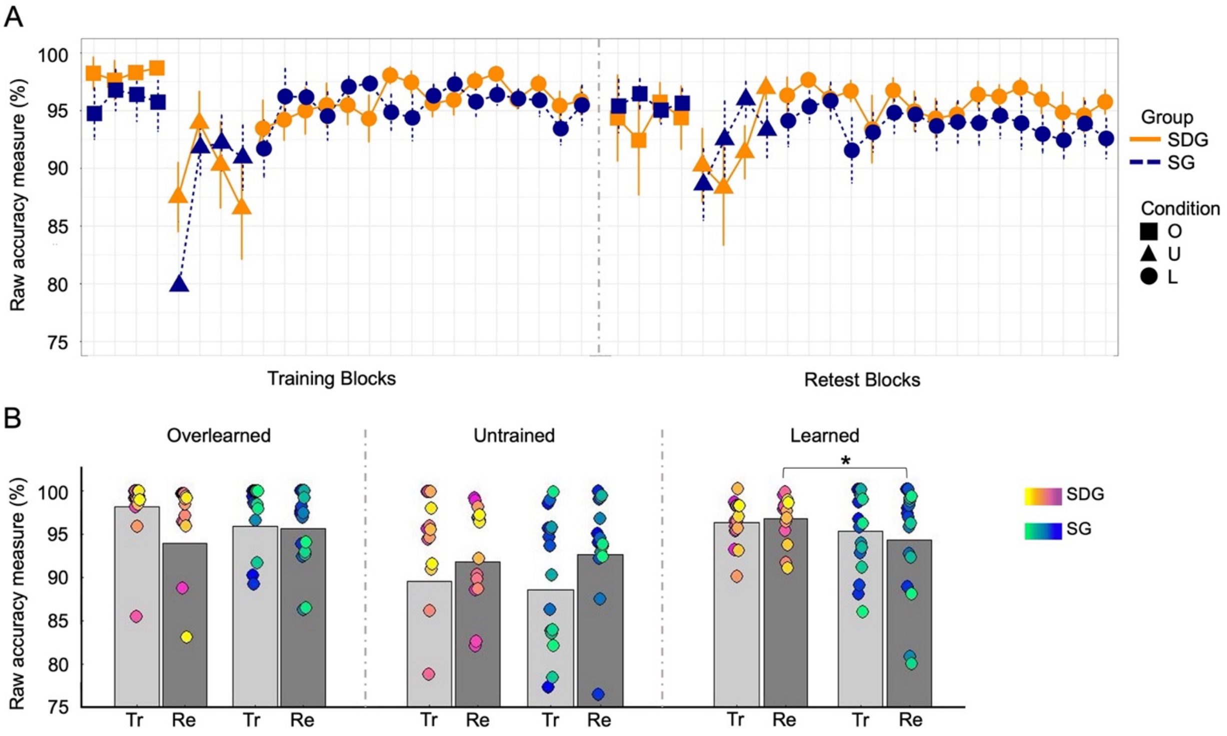


***Supplementary figure 1. Behavioral results (accuracy). A***: *Performance accuracy (proportion of corrects chunks) for each block of each condition during training and retest sessions for participants in the sleep group (SG, blue) and the sleep-deprived group (SDG, orange). Error bars represent SEM.* ***B:*** *Average performance across the last 4 blocks of training (Tr) and the first 4 blocks of retest (Re) for each condition in both groups (SDG, spring color bar; SG, winter color bar). Circles indicate individual participants. Significant (*) difference in performance between groups during the retest session for the L condition.*

*Supplementary figure 2*

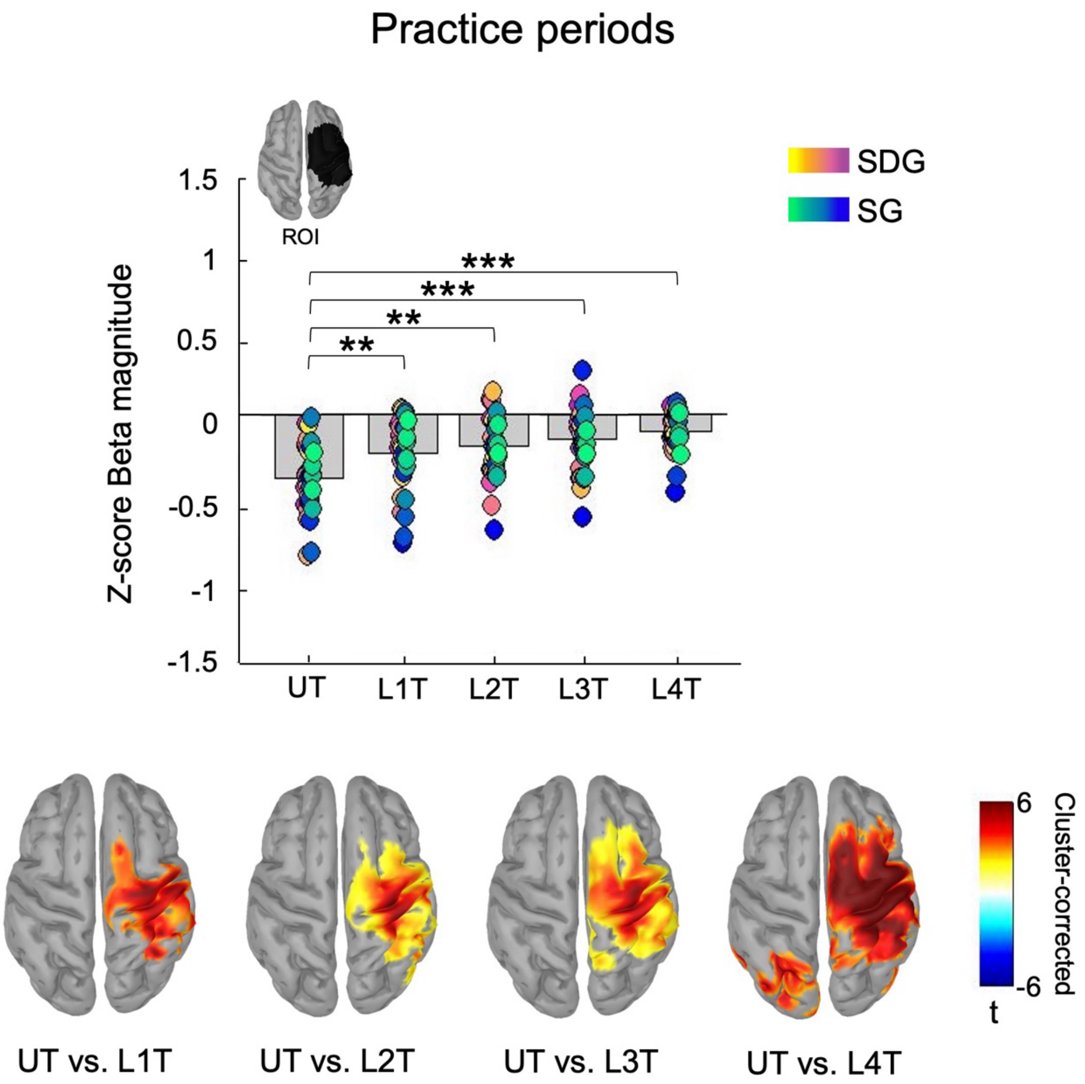


***Supplementary figure 2. Beta oscillations associated with task practice are modulated by learning during initial training.*** *Bar plot represents beta magnitude (Z-score) in motor regions (in the practice ROI) of all participants for the mean of 4 blocks for the UT, L1T, L2T, L3T and L4T conditions during the training session with colored circles indicating individual participants. Source maps represent significant contrasts (cluster corrected) performed at the whole brain level displayed on MNI-152 cortical mesh provided in SPM12 at a significance level* $\alpha$ *of .05. Results show a progressive decrease of beta desynchronization in right motor regions throughout practice, as compared to the UT condition.*

*Supplementary figure 3*

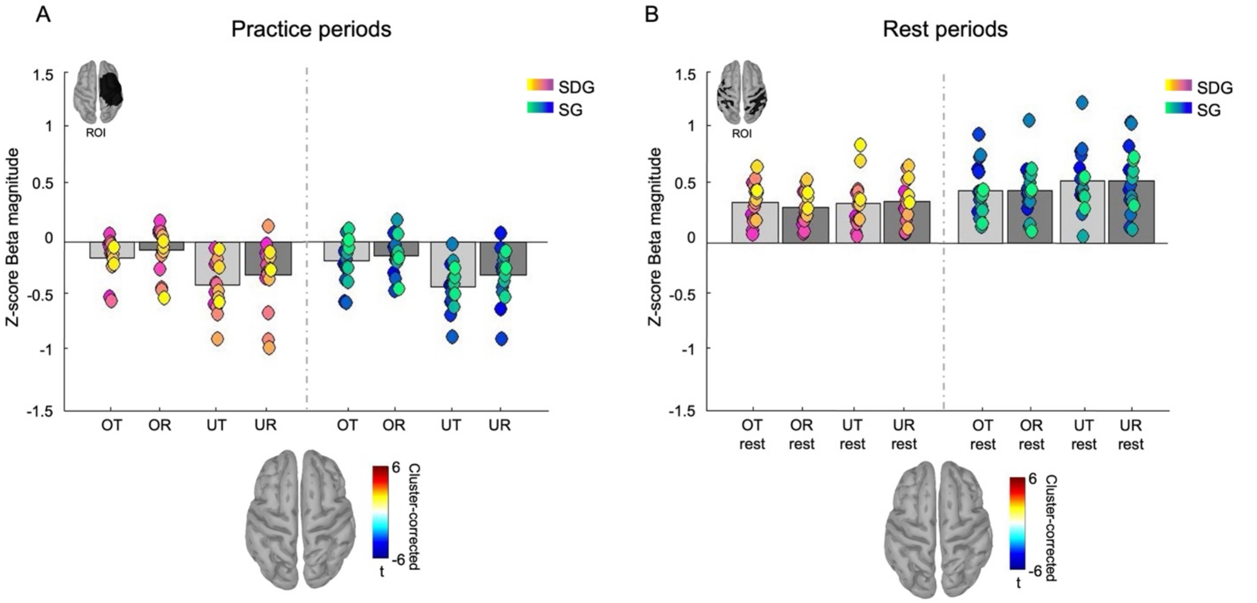


***Supplementary figure 3. Beta oscillations associated with task and inter-practice rest periods for the O and U conditions during the retest session in the SG and SDG. A:*** *Bar plot represents beta magnitude (Z-score) in motor regions (practice ROI) of all participants for O and U conditions during retest session separated by groups with colored circle indicating individual participants. Source maps represent significant contrasts (cluster corrected) performed at the whole brain level displayed on MNI-152 cortical mesh provided in SPM12 at a significance level $\alpha$ of .05. Beta oscillations during task did not differ between conditions and groups.* ***B:*** *Bar plot represents beta magnitude (Z-score) in parietal and motor regions (rest ROI) of all participants for rest periods following O and U conditions during retest session separated by groups with colored circles indicating individual participants. Source maps represent significant contrasts (cluster corrected) performed at the whole brain level displayed on MNI-152 cortical mesh provided in SPM12 at a significance level $\alpha$ of .05. Beta oscillations during rest periods did not differ between conditions and groups.*
